# Exploring the Phaeosphere: characterizing the microbiomes of *Phaeocystis antarctica* colonies from the coastal Southern Ocean and laboratory culture

**DOI:** 10.1101/2024.09.10.612332

**Authors:** Margaret Mars Brisbin, McCaela Acord, Rachel Davitt, Shavonna Bent, Benjamin A.S. Van Mooy, Eliott Flaum, Andreas Norlin, Jessica Turner, Arianna Krinos, Harriet Alexander, Mak Saito

**Affiliations:** Department of Marine Chemistry and Geochemistry, Woods Hole Oceanographic Institution (WHOI); University of South Florida College of Marine Science; Massachusetts Maritime Academy; Department of Marine and Coastal Sciences, Rutgers University; MIT-WHOI Joint Program in Oceanography / Applied Ocean Science and Engineering; Graduate Program in Biophysics, Stanford University; Department of Marine Sciences, University of Connecticut; Department of Biology, Woods Hole Oceanographic Institution

## Abstract

Interactions between phytoplankton and bacteria play critical roles in shaping marine ecosystems. However, the intricate relationships within these communities—particularly in extreme and rapidly changing environments like the coastal Southern Ocean—remain poorly understood. Here, we apply targeted methods to directly characterize the microbiomes of individual colonies of *Phaeocystis antarctica*, a keystone phytoplankton species in the Southern Ocean, for the first time. We show that colony microbiomes are consistent in distinct geographic locations at approximately the same time, but shift significantly after a year of laboratory culture. The bacterial orders Alteromonadales, Oceanospirillales, and Sphingomonadales dominated the microbiomes of all field-collected colonies, whereas Caulobacterales, Cellvibrionales, and Rhodobacterales dominated colony microbiomes after culturing. Notably, the most abundant genera in field-collected colony microbiomes, the psychrophiles *Paraglaciecola* and *Colwellia,* were lost in culture. The shift in microbiome structure emphasizes the importance of field-based studies to capture the complexity of microbial interactions, especially for species from polar environments that are difficult to replicate in laboratory conditions. Furthermore, the relative abundances of bacterial taxa comprising the majority of field-collected colony microbiomes—e.g., *Paraglaciecola sp.* (Alteromonadales) and Nitrincolaceae (Oceanospirillales)—were strongly associated with *Phaeocystis* abundance in surface waters, highlighting their potential roles in bloom dynamics and carbon cycling. This research provides valuable insights into the ecological significance of prokaryotic interactions with a key phytoplankton species and underscores the necessity of considering these dynamics in the context of climate-driven shifts in marine ecosystems.

## Introduction

Interactions between phytoplankton and bacteria form the basis of sophisticated and specialized relationships that influence large-scale ecosystem processes, such as primary production, nutrient cycling, and carbon export (Seymour et al. 2017). Primary production in the coastal Southern Ocean is impacted by the interactions between bacteria with key roles in ecosystem function (Kim et al. 2022) and specific phytoplankton groups (e.g., diatoms in the genera *Fragilariopsis* and *Pseudonitzschia*; (Bertrand et al. 2015)). The coastal Southern Ocean along the West Antarctic Peninsula is an exceptionally productive ecosystem with productivity driven by high-biomass phytoplankton blooms in the Austral spring and summer (Arrigo et al. 1998). The haptophyte alga *Phaeocystis antarctica* is a dominant early contributor to spring phytoplankton blooms and has a unique multi-morphic life history with implications for how carbon moves through the food web (DiTullio et al. 2000). *P. antarctica* has a free-living flagellate stage and a colonial stage, where thousands of cells coexist in a self-secreted, spherical mucilaginous colony (Schoemann et al. 2005). The colonial stage is dominant during *Phaeocystis* blooms (Smith and Trimborn 2024), is avoided by zooplankton grazers (Ryderheim et al. 2022), and contributes to organic carbon export through sinking (DiTullio et al. 2000, Nissen and Vogt 2021, Smith et al. 2021). The colony structure provides a unique interface for bacterial interactions that could contribute to *P. antarctica*’s ecological success. While *P. antarctica* blooms have been shown to alter the bacterial community in the water column (Delmont et al. 2014), the *P. antarctica* colony microbiome has not been directly investigated.

Evidence from other species suggests integral roles for the microbiome of *Phaeocystis* colonies. For example, the colonies of a closely related *Phaeocystis* species (*P. globosa*) host highly specific bacterial communities that can relieve B-vitamin limitation (Mars Brisbin et al. 2022). Bacteria isolated from the surface of the Southern Ocean diatom *Pseudo-nitzschia subcurvata* included a vitamin-B_12_ synthesizing *Sulfitobacter sp.* (Rhodobacterales) that boosted *P. subcurvata* growth rate and rescued it from B_12_ limitation (Andrew et al. 2022). Other bacteria isolated from *P. subcurvata* did not affect growth rate, but extended survival in stationary phase (Andrew et al. 2022). Earth system modeling predicts that particle-attached bacteria—like those associated with *Phaeocystis* colonies and other phytoplankton—will become more abundant in the Southern Ocean despite bacterial biomass in the oceans decreasing globally as mean sea surface temperature increases (Kim et al. 2023). These results highlight the relevance of particle and phytoplankton-associated bacteria under future climate scenarios, especially in the Southern Ocean (Kim et al. 2023). As the West Antarctic Peninsula is experiencing faster-than-average climate change (Schofield et al. 2010) and temperature-related shifts in phytoplankton communities (Schofield et al. 2017), fully characterizing interactions that could influence the abundance, productivity, and fate of keystone phytoplankton species, such as *P. antarctica,* is paramount.

The environmental conditions that organisms experience in the field are not perfectly emulated in laboratories in general, but especially in more extreme environments like the coastal Southern Ocean, where sea surface temperatures are lower than can be maintained by typical algal incubators and sea ice is an important physical feature. Thus, culture biases (the selection for or against certain organisms) in microbiome composition may be especially pronounced for organisms isolated from relatively extreme environments. The direct and immediate assessment of microbiome composition from field-derived isolates may better portray the bacterial community that exists in situ rather than studying the microbiomes of cultured isolates, but field microbiomes are often infeasible to analyze due to limited field access or challenges in identifying and isolating small phytoplankton cells at sea. Consequently, many studies characterizing phytoplankton-associated bacteria rely on accessible and easily manipulated cultured phytoplankton strains (Kuhlisch et al. 2024, Martínez-Pérez et al. 2024). However, the abundant large colonies comprising high-biomass *P. antarctica* blooms (Schoemann et al. 2005) enable direct and immediate microbiome assessment. Here, we isolated *P. antarctica* colonies from the surface of the Southern Ocean and immediately processed them for microbiome analysis. These microbiomes thus contained bacterial communities physically associated with *P. antarctica* colonies in natural bloom conditions. This approach is novel compared to previous studies that have used co-occurrence in bulk samples to identify bacteria associated with *Phaeocystis* (Delmont et al. 2014) or have evaluated individual-colony microbiomes in cultured strains (Mars Brisbin et al. 2022). Identifying the bacteria physically associated with *P. antarctica* colonies allows associations between *Phaeocystis* and bacteria to be authentically assessed in bulk samples. Using higher-resolution environmental samples in space and time in conjunction with direct and immediate validation of bacterial microbiome constituents can then enable identification and interpretation of which bacterial taxa have key roles in *Phaeocystis* blooms from formation to termination.

## Methods

Sampling was performed aboard the R/V *Nathanial B. Palmer* as part of the Palmer Long Term Ecological Research (PAL LTER) program’s annual cruise along the West Antarctic Peninsula completed from November 22 to December 17 during the Austral spring of 2021. Surface water was continuously sampled through the underway system and chlorophyll-*a* was monitored by a WET Lab ECO-FL Fluorometer, revealing especially high chlorophyll at offshore stations along the shelf-break (Figure 1). A Niskin bottle rosette was deployed at all PAL LTER grid stations, allowing for *P. antarctica* colonies to be non-destructively sampled from surface water at high-chlorophyll stations 400.200 and 600.200 (Figure 1; ∼108 nautical miles apart). Individual *Phaeocystis antarctica* colonies were isolated by transferring surface water (∼5m depth) from the bottom of Niskin bottles to glass plankton jars to prevent colony breakage from passing through the Niskin spout. Colonies were transferred to Petri dishes by pipette and observed by light microscopy to target intact colonies of similar size (diameters 86–161 µm, mean=125 µm; Supplemental Figure 1). Individual colonies were rinsed three times with 0.2 µm-filtered seawater before being transferred to an Axygen Maxymum Recovery 0.5 ml PCR tube with <10 µl of filtered seawater carried over. For DNA extraction, 30 µl of a 10% mass-to-volume Chelex 100 resin bead slurry in PCR-grade water was added to each colony, vortexed for 10 seconds, heated to 96°C for 20 minutes, vortexed again for 10 seconds, centrifuged briefly and transferred to ice. Polymerase chain reactions amplifying the V3–V4 region of the 16S rRNA gene were immediately performed on the supernatant from Chelex extractions using the Bento Lab Portable PCR Workstation. PCR conditions followed Illumina’s “16S Metagenomic Sequencing Library Preparation” protocol with no modifications. PCR success was checked by gel electrophoresis and successful reaction products were stored at –20°C before being hand-carried on ice back to the lab.

**Figure 1.**
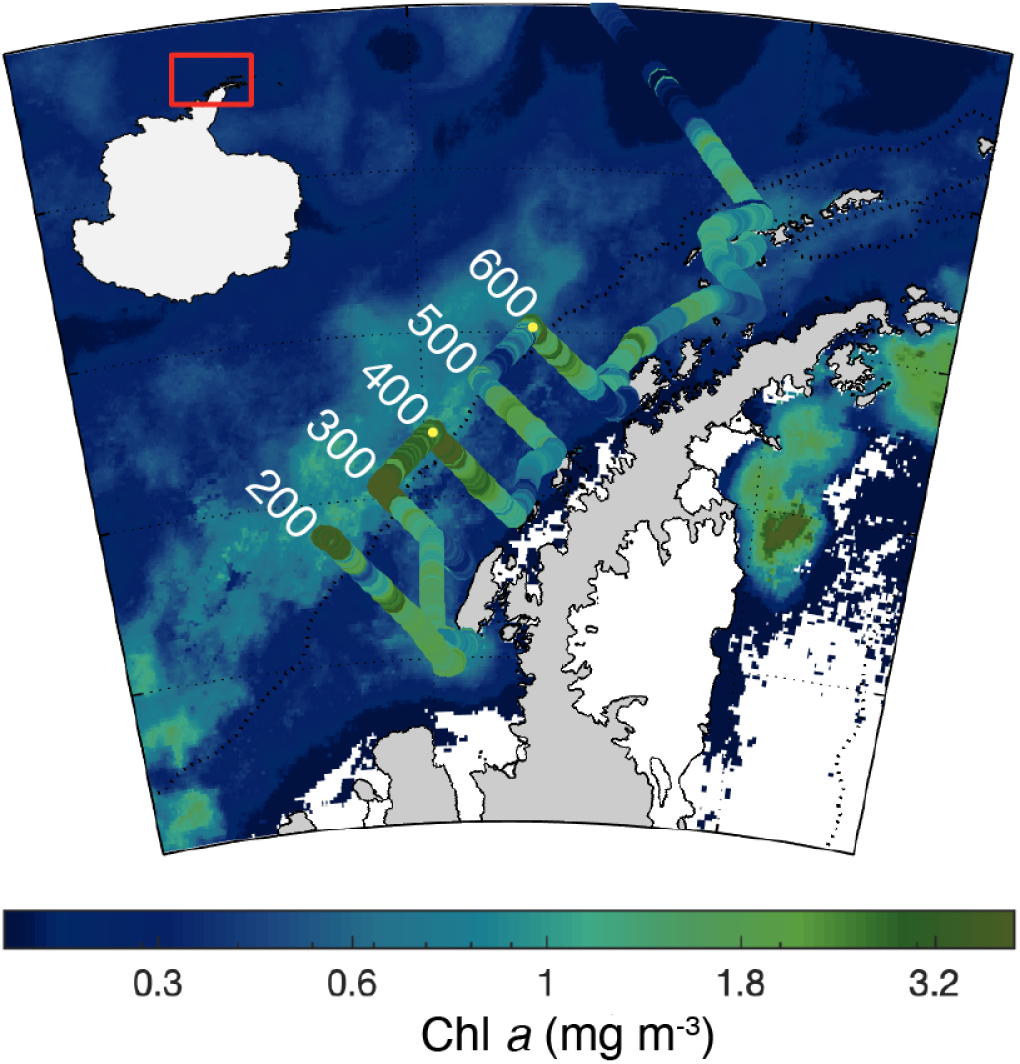
Chlorophyll-*a* concentration in surface seawater near the West Antarctic Peninsula during the sampling campaign. Satellite sea-surface chlorophyll-*a* was determined from CMEMS GlobColour Level-3 with the mean values from Nov 22–Dec 17, 2021 displayed (Garnesson et al. 2019). Underway chlorophyll-*a* measurements are overlaid on the satellite data using the same color scale. The study region is indicated with a red box on the inset map of Antarctica. A large offshore phytoplankton bloom is visible in the satellite data and corroborated by the underway measurements taken at the seaward ends of the Palmer Long Term Ecological Research program (PAL LTER) grid lines (labeled 200, 300, 400, 500, and 600 in white on the figure). Yellow points at the seaward ends of the 400 and 600 lines are stations 400.200 and 600.200, respectively, where *P. antarctica* colonies were collected for microbiome analysis.

To assess the effect of culture conditions on the *P. antarctica* colony microbiome composition, a unialgal culture was established by isolating a single colony from LTER grid station 400.200, rinsing it three times with sterile seawater, and transferring it to 35 mL of L1–Si media prepared with 0.2-µm filtered seawater (Guillard and Hargraves 1993) in a 50 mL plastic culture flask. At sea, the culture was maintained in a Percival low-temperature algal growth chamber with LED lighting, set to mimic ambient light and temperature conditions (–1°C, 18h: 6h light: dark cycle). The culture was hand-carried on ice back to the lab, where it was transferred to a walk-in cold chamber set to 4°C with 24-hour light from cool white fluorescent bulbs (recorded temperature range: 4–8°C). The culture was maintained in 100 mL of media (L1–Si in 34 PSU artificial seawater prepared from Instant Ocean) in a plastic culture flask with 75% of the media replaced approximately once a month. After about a year of maintenance (in January 2023), individual colonies (n=14) were isolated from the culture and processed for single-colony microbiome analysis following the same methods as the field-isolated colonies described above.

To evaluate the influence of *Phaeocystis* abundance on the bacterial community composition in surface waters, we performed whole-community 16S rRNA metabarcoding (cDNA) with samples collected from the sea surface in the LTER sampling area. Two liters of surface water collected by Niskin bottle at grid stations were filtered through 47 mm-diameter 0.2-µm-pore-size filters. To increase the spatial resolution of environmental sampling, seawater from the ship’s underway system was also filtered during selected transects between grid stations. Seawater from the underway system was screened through 51 µm-mesh and filtered on 3.0 µm-pore-size 142 mm-diameter filters using positive pressure for one hour at a time. The GPS coordinates when filtering began and ended for each underway sample were recorded (samples integrated 3–23 nautical miles, mean=10.25 nautical miles), along with the volume of water filtered (5.3–18 L, mean=11 L). All filters were flash-frozen in liquid nitrogen, stored at –80°C onboard, and transferred to coolers with dry ice for shipping to the lab. Upon arrival, RNA was extracted from the 0.2-µm-pore-size filters (Niskin samples from grid stations) and one-quarter sections of the 3.0 µm-pore-size filters (underway samples) using the Qiagen RNeasy Mini Kit with additional mechanical cell lysis using Zymo Scientific Bashing Beads (0.1 & 0.5 mm diameter). RNA was extracted from these samples to target the active portion of the environmental communities as much as possible. RNA was reverse transcribed to cDNA using the Applied Biosystems High-Capacity RNA-to-cDNA kit before proceeding with PCRs to amplify the V3–V4 region of 16S rRNA.

PCR products from field-collected colonies, cultured colonies, and filtered seawater samples were sequenced with 300 x 300-bp v3 paired-end chemistry on the Illumina MiSeq Platform, with sequencing completed on two flow cells one year apart, but at the same sequencing facility following the same methods. Sequences from the two sequencing runs were processed separately with the DADA2 denoising algorithm (Callahan et al. 2016) within the QIIME 2 framework (Bolyen et al. 2019), which performs quality filtering, chimera and singleton removal, and amplicon sequence variant (ASV) identification. Following denoising, the results from the two runs were merged and the ASVs were clustered at 99% identity. Taxonomy was assigned to all representative sequences using a naive Bayes classifier trained on the SILVA 99% 16S sequence database (v138; Quast et al. 2013) using the QIIME 2 feature-classifier plug-in (Bokulich et al. 2018). Sequences classified as chloroplasts with the SILVA classifier were transferred to a new fasta file and reclassified with a naive Bayes classifier trained on the PhytoREF database (Decelle et al. 2015). Further analysis was completed in the R statistical environment (R Core Team 2018) using the packages phyloseq (v1.34; McMurdie and Holmes 2013), CoDaSeq (v0.99.6; Gloor et al. 2017), and vegan (v2.5–7; Oksanen et al. 2019).

## Results

Collectively, the field-collected colonies hosted 241 unique bacterial ASVs in their microbiomes, with 28–67 bacterial ASVs detected per colony (mean=44, n=26; Figure 2A). Cultured-colony microbiomes were less diverse, with 74 unique ASVs detected across all cultured-colony microbiomes and 18–31 ASVs detected per colony (mean=25, n=14; Figure 2A). While a large proportion of sequences from single-colony samples were *Phaeocystis sp.* chloroplast sequences (52–96%, mean=72.8%; Supplemental Figure 2), rarefaction analysis showed that bacterial ASV richness saturated within the total number of bacterial sequences for each sample (Supplemental Figure 3). Microbiome community compositions were consistent among field-collected colonies as well as among cultured colonies, but were significantly different between field-collected and cultured colonies (PERMANOVA, 999 permutations, *F*=44.209, *R*^2^=0.54, *p*=0.001; Figure 2B, Supplemental Figure 4). Alteromonadales was the most abundant bacterial order in the field-collected colony microbiomes (Figure 2B), with the majority of Alteromonadales ASVs belonging to the genera *Paraglaciecola* and *Colwellia* (Supplemental Figures 5, 6). Notably, a single *Paraglaciecola sp.* ASV comprised a large proportion of almost all field-collected colony microbiomes (mean=32%, sd=13%, range=11–57%, when excluding one sample with only 1% of this ASV; Supplemental Figure 5). In contrast, the orders Caulobacteriales, Cellvibrionales, and Rhodobacterales made up the majority of the cultured-colony microbiomes (Figure 2). Within Caulobacteriales, two ASVs in the genus *Algimonas* together contributed 6–30% of cultured-colony microbiome sequences (mean=18.2%). Two dominant ASVs made up the majority of Cellvibrionales sequences, with one belonging to the genus *Sinobacterium* (mean=21%, range=2–39%) and the other only classified to family-level (Family: Cellvibrionaceae; mean=30%, range=6–60%). A single *Sulfitobacter* ASV contributed most of the Rhodobacterales sequences in cultured-colony microbiomes (mean=22%, range=12–31%; Supplemental Figure 7). Overall, the cultured-colony microbiomes had reduced diversity (ASV richness) and lost most *Paraglaciecola* and all *Colwellia* members along with Oceanospirillales (family: Nitrincolaceae) and Sphingomonadales (family: Erythrobacteraceae) members. In the absence of these major constituents of field-collected colony microbiomes, the Caluobacteriales, Cellvibrionales, and Rhodobacterales appear to thrive in culture conditions.

**Figure 2.**
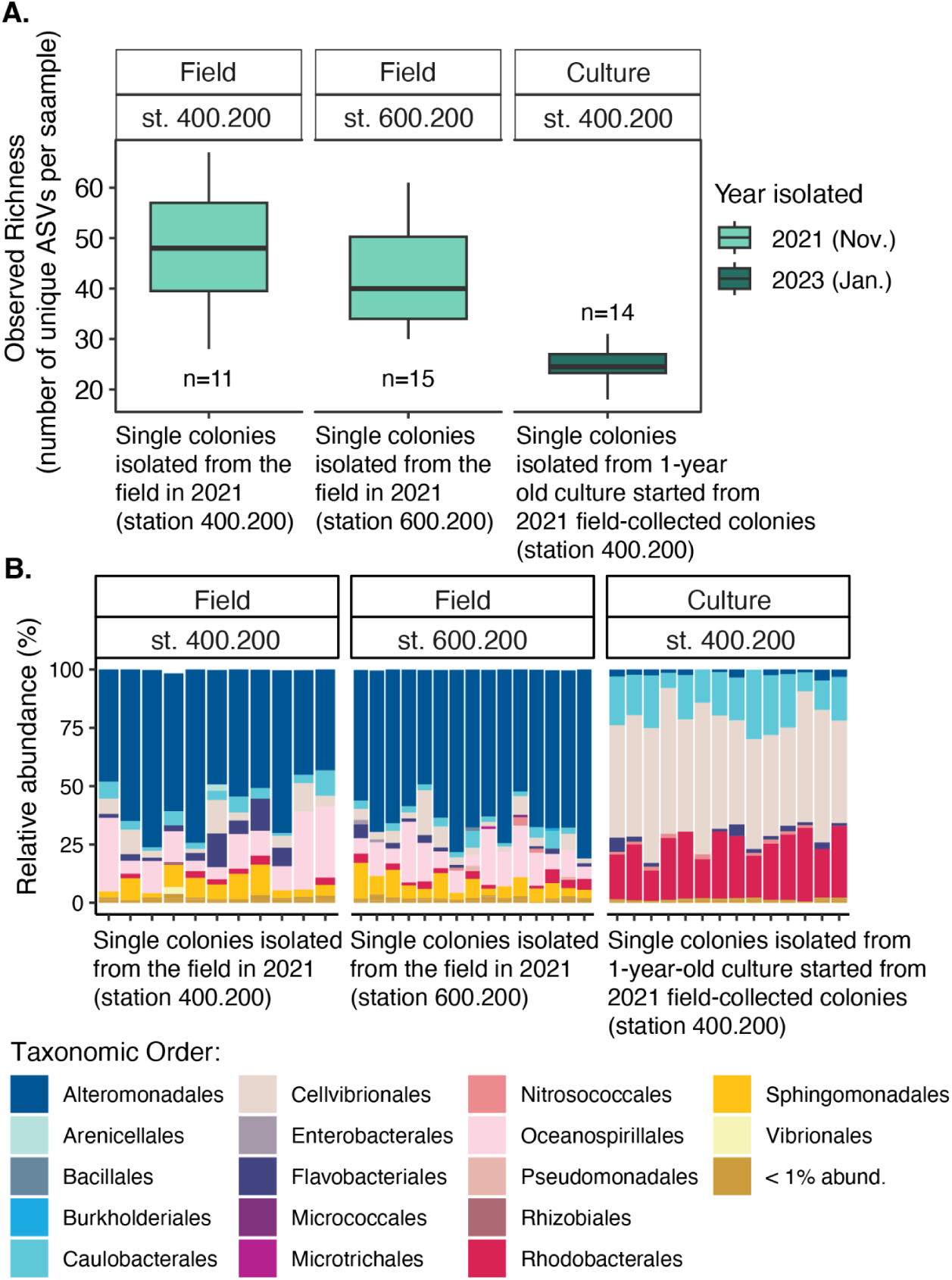
Diversity and community composition of bacterial communities associated with single *Phaeocystis antarctica* colonies isolated directly from the field or after a year in culture. **(A)** The observed richness (i.e., number of unique amplicon sequence variants) in each colony microbiome. The solid back lines inside boxes represent the median, boxes represent the interquartile range between the 1^st^ and 3^rd^ quartiles, and whiskers denote the range. Sample size (n) is indicated above or below each box and represents the number of individual colonies analyzed. **(B)** Relative abundance of bacterial orders in each colony microbiome (each column represents a single colony).

A clear geographical pattern in *P. antarctica* relative abundance was apparent in the chloroplast 16S rRNA sequences from environmental water samples (Niskin and underway samples; Figure 3). The abundance of *Phaeocystis* relative to other phytoplankton groups increased from the mid-shelf to off-shelf stations, with ∼50% of chloroplast 16S rRNA sequences belonging to *Phaeocystis* along the shelf break (Figure 3A–B). Samples collected closer to shore included larger proportions of prasinophytes and silicoflagellates (families: Prasinophyceae and Dictyochophyceae; Figure 3A–B). The bacterioplankton community showed a similar geographic pattern, with communities shifting from higher diversity closer to shore to lower diversity at the shelf break (Figure 3C–D). Flavobacteriales were more abundant at the shelf-break and at stations with higher *Phaeocystis* abundance, whereas SAR11 was more abundant closer to shore and at stations with lower *Phaeocystis* abundance.

**Figure 3.**
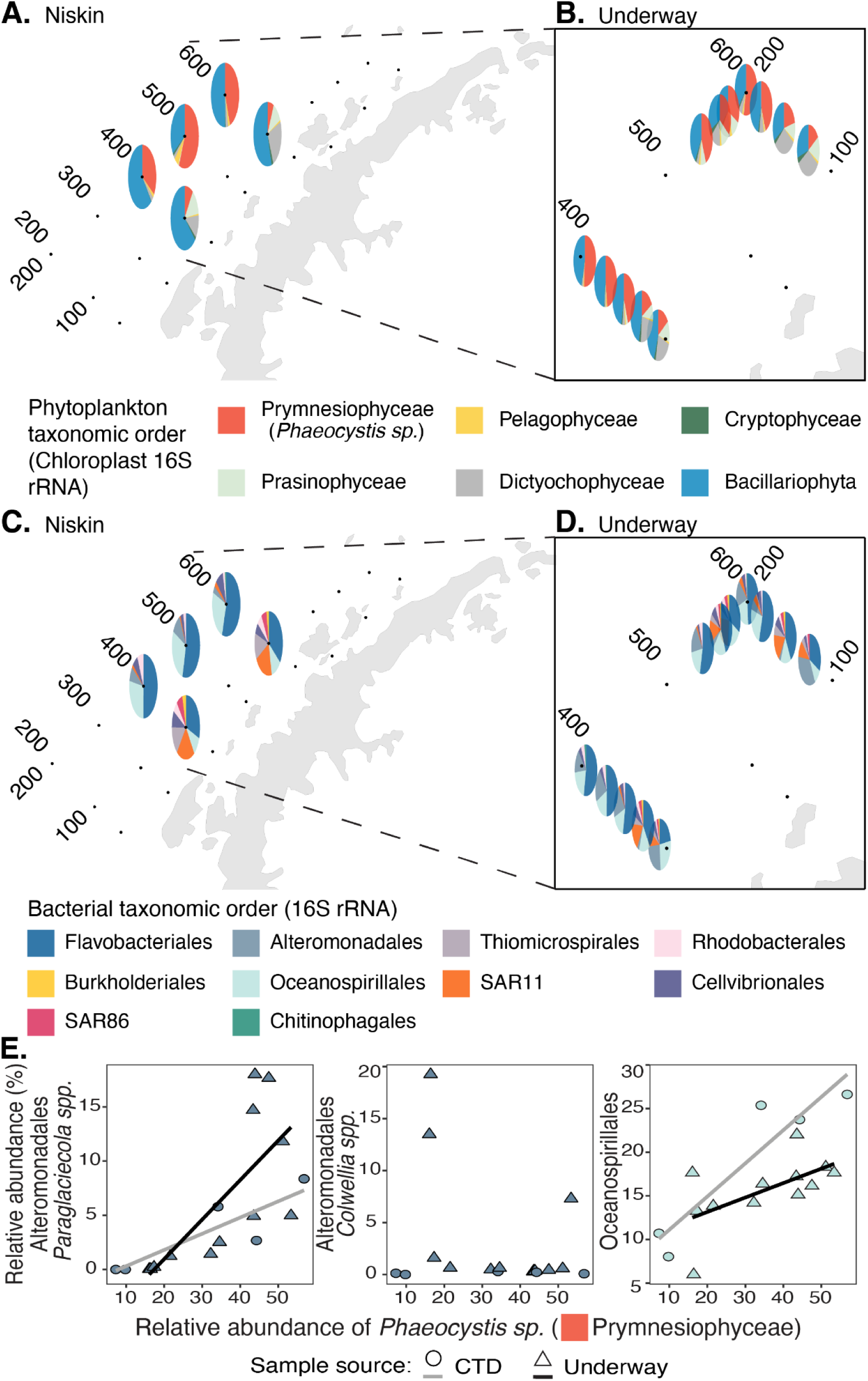
Relative abundances of major phytoplankton and bacterial taxonomic orders and the relationship between abundances of major components of native *P. antarctica* colony microbiomes and *P. antarctica* in the Palmer LTER grid region. **(A–B)** Relative abundances of major phytoplankton orders in Niskin and underway samples based on chloroplast 16S rRNA sequences are denoted by pie charts arranged to match the geographic location where the samples were collected. **(C–D)** Relative abundance of major bacterial orders in Niskin and underway samples based on 16S rRNA sequences. Palmer LTER grid stations are represented by black points and the grid lines are labeled. Data were plotted with the R packages ggspatial and scatterpie. **(E)** Relationship between the relative abundance of dominant Alteromonadales ASVs (*Paraglaciecola spp.* and *Colwellia spp.*) and Oceanospirillales ASVs in field-collected colony microbiomes with *Phaeocystis* chloroplast 16S rRNA in environmental samples (surface seawater) from the Palmer LTER grid. For this analysis, the environmental dataset was subset to only include ASVs that contributed majority proportions of field-collected colony microbiomes. Linear regression lines are plotted when correlations between bacteria and *Phaeocystis* abundances were statistically significant (*p* ≤ 0.05).

The environmental bacterial communities were further interrogated to investigate how the relative abundance of bacterial ASVs directly associated with *Phaeocystis* colonies changed with the relative abundance of *Phaeocystis* in surface seawater. Alteromonadales made up the majority of field-collected colony microbiomes, with the genera *Paraglaciecola* and *Colwellia* contributing most of these sequences (Figure 2, Supplemental Figures 5, 6). The relative abundance of *Paraglaciecola* ASVs significantly correlated with the relative abundance of *Phaeocystis* in both the Niskin (*R*^2^=0.77, *p*=0.05) and underway samples (*R*^2^=0.52, *p*=0.008), but the *Colwellia* ASVs were generally low-abundance in environmental samples and did not correlate to *Phaeocystis* abundance (Figure 3E). Twenty-eight Oceanospirillales ASVs were detected in field-collected colony microbiomes, with two of these prominently prevalent and abundant across microbiomes (Supplemental Figures 8, 9). Of the 28 colony-associated Oceanospirillales ASVs, nine were also detected in the environmental samples, including the two that were most abundant in colony microbiomes. Summed relative abundances of Oceanospirillales ASVs detected in both microbiome and environmental samples significantly correlated to the relative abundance of *Phaeocystis* in both Niskin (*R*^2^=0.87, *p*=0.02) and underway samples (*R*^2^=0.36, *p*=0.04; Figure 3E). The most abundant Oceanospirillales ASVs belonged to the family Nitrincolaceae but were generally not able to be classified further (Supplemental Figures 9, 10). Sphingomonadales was the third most abundant order in field-collected colony microbiomes, but Sphingomonadales ASVs were not detected in most environmental samples (max. rel. abundance in any sample=0.04%) and the Sphingomonadales ASVs found in colony microbiomes were not detected in any environmental samples.

## Discussion

By directly evaluating microbiomes of *P. antarctica* colonies from the field, we demonstrated that *P. antarctica* colonies host consistent microbiomes even when sampled from geographically distinct locations (∼108 nautical miles apart) within a single season. This result is consistent with previous work showing that *Phaeocystis globosa* colonies hosted similar microbiome communities regardless of where the culture originated (Mars Brisbin et al. 2022). However, when *P. antarctica* was cultured and maintained in a walk-in cold chamber with 24-hour fluorescent lighting for one year, the microbiome diversity was halved and the overall composition shifted from being mainly composed of Alteromonadales, Oceanospirillales, and Sphingomonadales to being dominated by Caulobacterales, Cellvibrionales, and Rhodobacterales (Figure 2). The major constituent of field-collected *P. antarctica* colony microbiomes was *Paraglaciecola sp.* (Alteromonadales), a psychrophile previously found in association with sea ice and ice algae (Vadillo Gonzalez et al. 2022). The second most abundant field-collected colony microbiome constituent, *Colwellia sp.*, is also psychrophilic (Methé et al. 2005, Zhang et al. 2017). These psychrophiles were lost or reduced in culture, probably due to temperature variation during transport and incubation at ∼4°C instead of –1°C. However, the precise loss point or cause can not be definitively identified because the cultured-colony microbiome was only evaluated at the end of the culture period. Rhodobacterales—a group often reported to associate with phytoplankton phycospheres (Beiralas et al. 2023, Roager et al. 2023)—were a dominant contributor to cultured-colony microbiomes but were a minority in field-collected colony microbiomes. Rhodobacterales may be especially suited to laboratory conditions typical of phytoplankton culture and, therefore, may increase in relative abundance during culture. These results highlight the importance of investigating microbial interactions in situ, especially when environmental conditions are difficult to reproduce, as is true of extreme polar environments.

To assess the significance of *P. antarctica*-associated bacteria in a broader environmental context, the relative abundances of ASVs belonging to major colony-associated taxonomic groups (Alteromonadales, Oceanospirillales, Sphingomonadales) were compared to the relative abundance of ASVs classified as *Phaeocystis* chloroplast in each environmental sample. *Paraglaciecola* (Alteromonadales) relative abundance was strongly correlated to *Phaeocystis* abundance, demonstrating that this dominant microbiome constituent was more abundant in surface seawater when *Phaeocystis* was more abundant (Figure 3). *Paraglaciecola spp.* are motile (Wang et al. 2020), particle-associated marine bacteria (Heins et al. 2021) with a diverse array of functional carbohydrate-active enzymes (CAZymes) that allow them to metabolize algae-specific complex polysaccharides, including agars, carrageenans, alginates, and fucoidans (Schultz-Johansen et al. 2018). Therefore, *Paraglaciecola* may be particularly suited to actively colonize *P. antarctica* colonies and utilize the abundant and diverse polysaccharides that comprise the *P. antarctica* colonial matrix and are excreted by *P. antarctica* cells (Alderkamp et al. 2007). Similar to *Paraglaciecola* ASVs, Oceanospirillales ASVs (family: Nitrincolaceae) from the colony microbiomes were more abundant at the sea surface when *Phaeocystis* was more abundant (Figure 3, Supplemental Figure 10). While less is known about their metabolic capabilities, Nitrincolacea abundance is correlated to phytoplankton biomass in polar waters (Liu et al. 2020, Wietz et al. 2021, Thiele et al. 2023) and Nitrincolacea are often first responders to increasing organic carbon availability at the start of phytoplankton blooms (Liu et al. 2020, Thiele et al. 2023).

In contrast to *Paraglaciecola* and Nitrincolaecea, *Colwellia spp.* (Alteromonadales) abundance was generally lower in surface water samples and uncorrelated with *Phaeocystis* abundance, despite being relatively abundant in all field-collected colony microbiomes (Figure 3). A previous study comparing bacterial communities inside and outside Southern Ocean *Phaeocystis* blooms found *Colwellia* abundance correlated with *Phaeocystis* abundance at depth, but not at the surface, leading the authors to hypothesize that *Colwellia* was active in degrading sinking *Phaeocystis* colony biomass (Delmont et al. 2014). Corroborating this hypothesis, another study found that *Colwellia spp.* were initially rare within bacterial communities on sinking particles collected in the Arctic, but became abundant after a ten-day incubation (Heins et al. 2021). In our present study, *Colwellia* was found to be prevalent across field-collected colony microbiomes by assessing the 16S rRNA gene (DNA), whereas the environmental communities in surface seawater were assessed with 16S rRNA (RNA) and *Colwellia* was barely detected in most of these samples. Sequences originating from RNA instead of DNA provide insight into activity in addition to abundance (Li et al. 2017), suggesting that the *Colwellia* in colony microbiomes could be relatively inactive. Colony-associated *Colwellia* may become more active in response to senescence-associated changes in the *Phaeocystis* colony microhabitat as *Phaeocystis* sinks and senesces at depth. Although additional sampling would be necessary to directly test this, phycosphere-associated bacteria have been shown to respond metabolically to algal senescence (Seyedsayamdost et al. 2011). *Colwellia* could also become more active and abundant in surface waters later in the season as the *Phaeocystis* bloom declines and colonies senesce before sinking. Ultimately, time-course single-colony, surface-water, and subsurface-water samples collected from the beginning of the bloom through bloom demise would be needed to fully evaluate the role of *Colwellia* in degrading *Phaeocystis* colony biomass.

SAR92 (Order: Cellvibrionales) was previously reported as the bacterial group most strongly correlated with *Phaeocystis* abundance in surface waters and was also found to be enriched in the particulate size fraction during *Phaeocystis antarctica* blooms (Delmont et al. 2014). A SAR92 ASV contributed 0–18% (mean=2.8%) of bacterial 16S sequences generated for field-collected colony microbiomes (absent in cultured-colony microbiomes), confirming this previously hypothesized association. The SAR92 ASV ranged in relative abundance from 0.6–8.2% (mean=3.5%) in our surface water samples, which is lower than the 30–40% of sequences from >3 µm samples reported in Delmont et al. (2014). The SAR92 ASV was significantly correlated with *Phaeocystis* abundance in our underway samples (*R*^2^=0.54, *p*=0.006)—which excluded the 0.2–3-µm-size fraction—but not the CTD samples (*R*^2^=0.3, *p*=0.34) that were unfractionated. SAR92 relative abundance increases during phytoplankton blooms, especially at high latitudes, suggesting it may generally respond to phytoplankton biomass and phytoplankton-derived organic matter (Wemheuer et al. 2015, Teeling et al. 2016, Liu et al. 2020, Xue et al. 2021). However, SAR92 can use dimethylsulfoniopropionate (DMSP) as its sole carbon source by enzymatically cleaving DMSP and releasing climatically active dimethyl sulfide (DMS) (He et al. 2023). When experimentally exposed to elevated DMSP, SAR92 upregulates expression of the DMSP cleavage pathway, demonstrating that DMSP is an important carbon source for these bacteria (He et al. 2023). As *Phaeocystis* is a prolific producer of DMSP (van Duyl et al. 1998, DiTullio et al. 2000, Vance et al. 2013), SAR92 may be especially poised to respond to *Phaeocystis* blooms and associate with *Phaeocystis* colonies. Moreover, the physical coupling of SAR92 bacteria and *P. antarctica* colonies may expedite DMS production and release to the atmosphere, influencing cloud formation, atmospheric processes, and climate feedbacks (Park et al. 2021).

Phytoplankton-bacteria interactions have the potential to drive bloom formation, decline, and succession, with consequences for the food web and carbon sequestration. *Phaeocystis* is an important contributor to primary production and carbon export in the Southern Ocean, but many aspects of its physiology–particularly drivers of its blooms–remain enigmatic. We showed that *Phaeocystis antarctica* colonies have consistent microbiomes when collected from two sites in the Palmer LTER grid. Most of the dominant field-collected colony-associated bacteria can move towards regions of interest, colonize particles, and metabolize polysaccharides and nitrogen compounds that are relatively unique to the *Phaeocystis* colonial matrix. Thus, bacteria associated with colonies likely benefit from the association through chemical currencies made available by *Phaeocystis* cells. However, it remains unclear as to how or if *Phaeocystis* benefits from hosting these specific microbiome communities. *Paraglaciecola* followed by *Colwellia* were the most prevalent microbiome constituents for field-collected colonies. All of the completed *Paraglaciecola* (n=3) and *Colwellia* (n=7) genomes that are annotated in the KEGG database (Kanehisa et al. 2023) have complete salvage and repair pathways for building vitamin B_12_ from precursor cobamides (Rodionov et al. 2003, Fang et al. 2017). A marine *Colwellia* isolate was shown to require B_12_ for growth, but was rescued from B_12_ limitation if cobamides were supplied. Moreover, when *Colwellia* was grown on cobamides, it shared B_12_ with two different B_12_-requiring diatoms, rescuing them from B_12_ limitation in experimental conditions. The *Colwellia* also released another B_12_ precursor, the lower ligand α-ribazole, that could be used by other bacteria to build B_12_ (Wienhausen et al. 2024). Thus, it is conceivable that microbiome bacteria, either individually or in concert, provide *P. antarctica* with B_12_, a service that is common among phytoplankton microbiomes (e.g., Durham et al. 2015, Cruz-López and Maske 2016) and that would give *P. antarctica* a competitive boost at the start of the spring bloom (Bertrand et al. 2007, Joy-Warren et al. 2022). Given that this study emphasizes the importance of studying microbial interactions in situ and cautions against evaluating microbial interactions in culture conditions (especially for polar species), culture experiments investigating the roles of *P. antarctica* colony microbiomes may not be representative. Instead, the direct sequencing of *Phaeocystis*—and other phytoplankton—microbiomes provides a road map for which bacteria to target in meta-omic analyses. For instance, future work may target the transcriptional activity of *Paraglaciecola* in the particulate size fraction of seawater samples from a *Phaeocystis* bloom to elucidate benefits bestowed by colony microbiomes.

The West Antarctic Peninsula region is undergoing rapid environmental change, with air and sea surface temperatures having increased ∼6°C and 1°C, respectively, since the mid-1900s (Meredith and King 2005). Warming has caused glacial retreat (Cook et al. 2016) and reduced sea ice coverage and annual duration in the region (Stammerjohn et al. 2021). With phytoplankton community composition and bloom biomass tightly coupled to seasonal sea-ice retreat and its impact on mix layer depth and nutrient availability, ongoing shifts in dominant phytoplankton bloom taxa (Nardelli et al. 2023) and phenology (i.e., timing of annual bloom events) are expected to continue (Turner et al. 2024). While the overall spring bloom onset is shifting later in the year, the bloom in the offshore region—where *P. antarctica* is more likely to be abundant (Joy-Warren et al. 2019)—is occurring slightly earlier (Turner et al. 2024). Increasing temperature and changing environmental conditions associated with altered bloom timing are likely to influence phytoplankton microbiome composition and function, especially if major microbiome constituents are psychrophilic. While the loss of major microbiome constituents in the culture conditions in this study (increased temperature, higher nutrient availability, extended photoperiod, and altered composition of DOM) suggests changing climate will influence microbiome diversity, increased temperature, and shortened photoperiod at bloom onset could also tip microbiome relationships to become more antagonistic (Giesler et al. 2023, Lin et al. 2024). As environmental conditions continue to shift with global climate change, it will be increasingly important to comprehensively characterize interactions between phytoplankton and their microbiomes and determine how changes to these relationships may propagate through the ecosystem.

## Acknowledgements

We thank the captain and crew of the R/V Nathaniel B. Palmer along with the scientists and scientific support staff associated with the PAL LTER program for their assistance in sample collection and underway measurements. This work was supported by NSF OPP-2224611 to BASVM. MMB was supported by the Simons Foundation through a Marine Microbial Ecology Postdoctoral Fellowship (award 874439). MA was supported through the Blue Economy Internship program at WHOI, which is supported in part by John and Shirley Farrington. We further thank Gretta Serres, Julie Huber, and Kama Thieler for their leadership roles in the Blue Economy Internship program. Sequencing for this project was completed by the Georgia Genomics and Bioinformatics Core (GGBC, UG Athens, GA, RRID:SCR_010994).

Authors have no conflicts of interest to declare.

## Data Availability

All sequencing data generated for this project are available from the NCBI SRA under accession PRJNA1136990. Intermediate data files and the code necessary to replicate analysis are available in a GitHub repository (https://github.com/MICOlab-USF/phaeosphere_AntarcticEdition) and as an interactive HTML document (https://micolab-usf.github.io/phaeosphere_AntarcticEdition/Phaeosphere_Analysis.html).

## Supplemental Figures

**Supplemental Figure 1.**
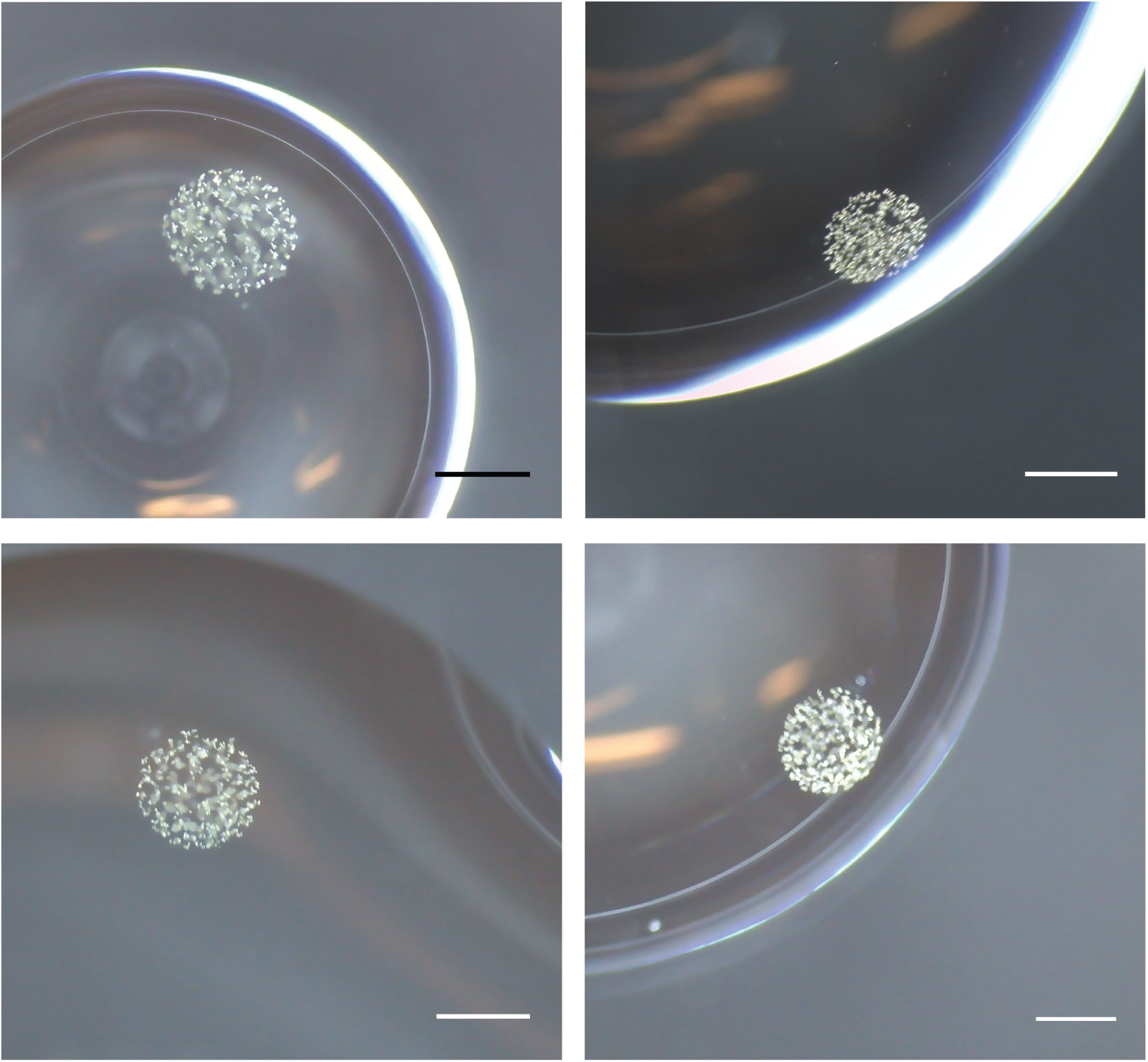
Representative *Phaeocystis antarctica* colonies isolated from station 600.200 in the PAL LTER grid and processed for microbiome analysis. Scale bars are 100 µm and images were taken at 200x magnification during the final rinse before transfer to PCR tubes for DNA extraction.

**Supplemental Figure 2.**
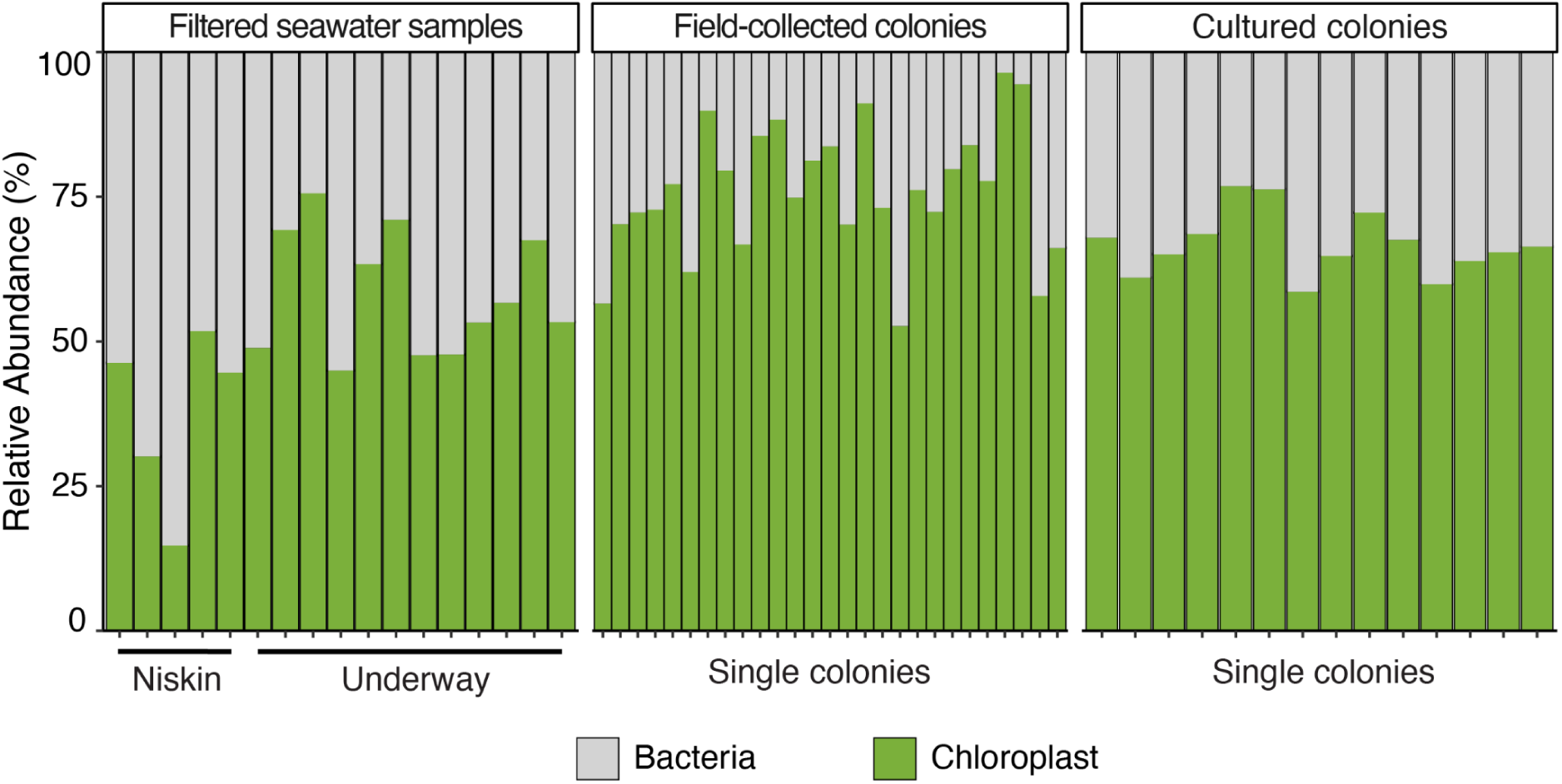
Relative abundance of bacterial and chloroplast 16S rRNA sequences in all samples. Bacterial and chloroplast sequences were split and analyzed separately. Single colony samples had 52–96% chloroplast sequences.

**Supplemental Figure 3.**
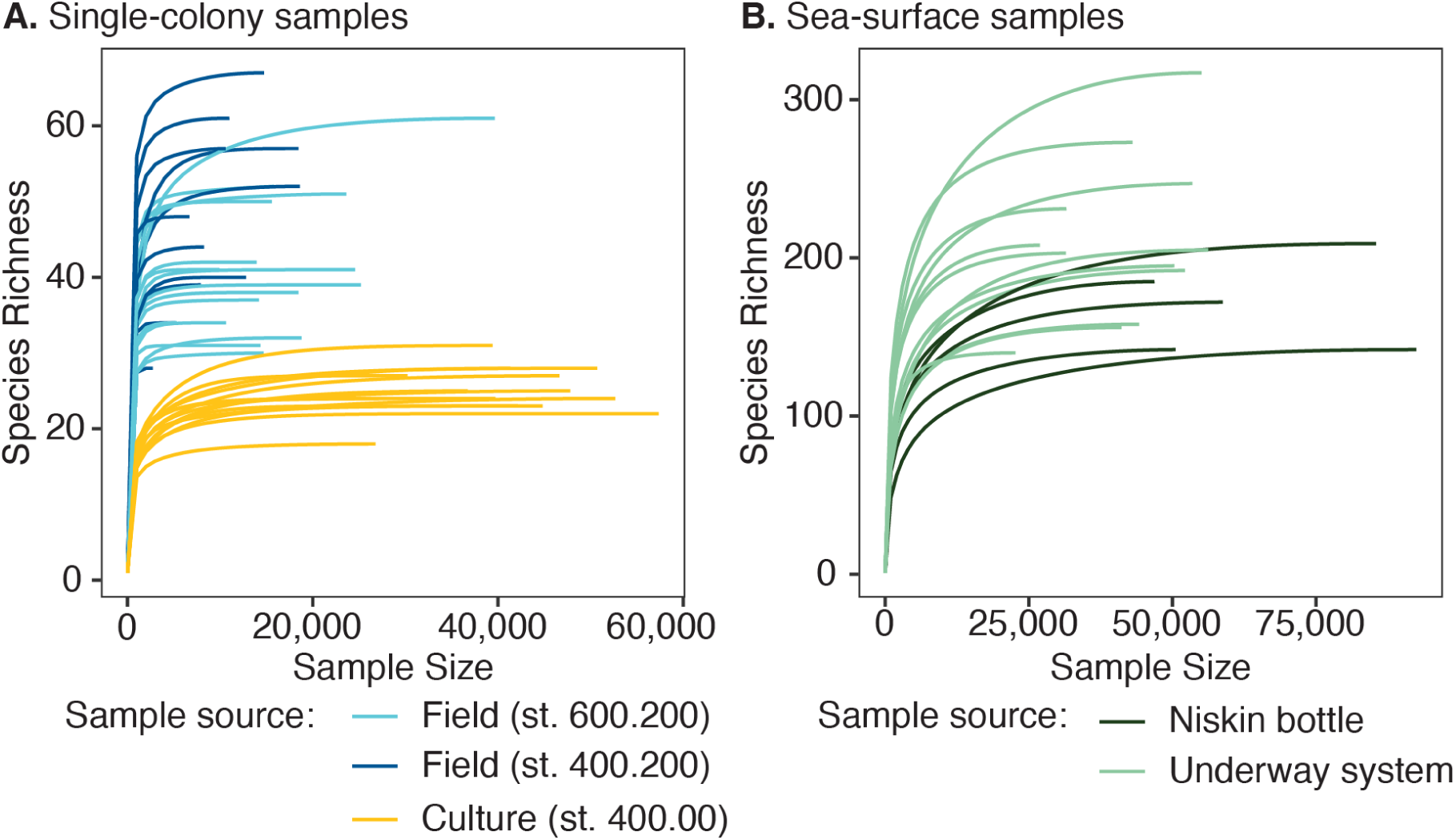
Rarefaction curves for bacterial sequences in single-colony and surface seawater samples. Rarefaction was performed with the R package phyloseq after chloroplast sequences were removed from the data and results were plotted with the ggrare function. All samples reached species (ASV) richness saturation within the total number of sequences generated for the sample.

**Supplemental Figure 4.**
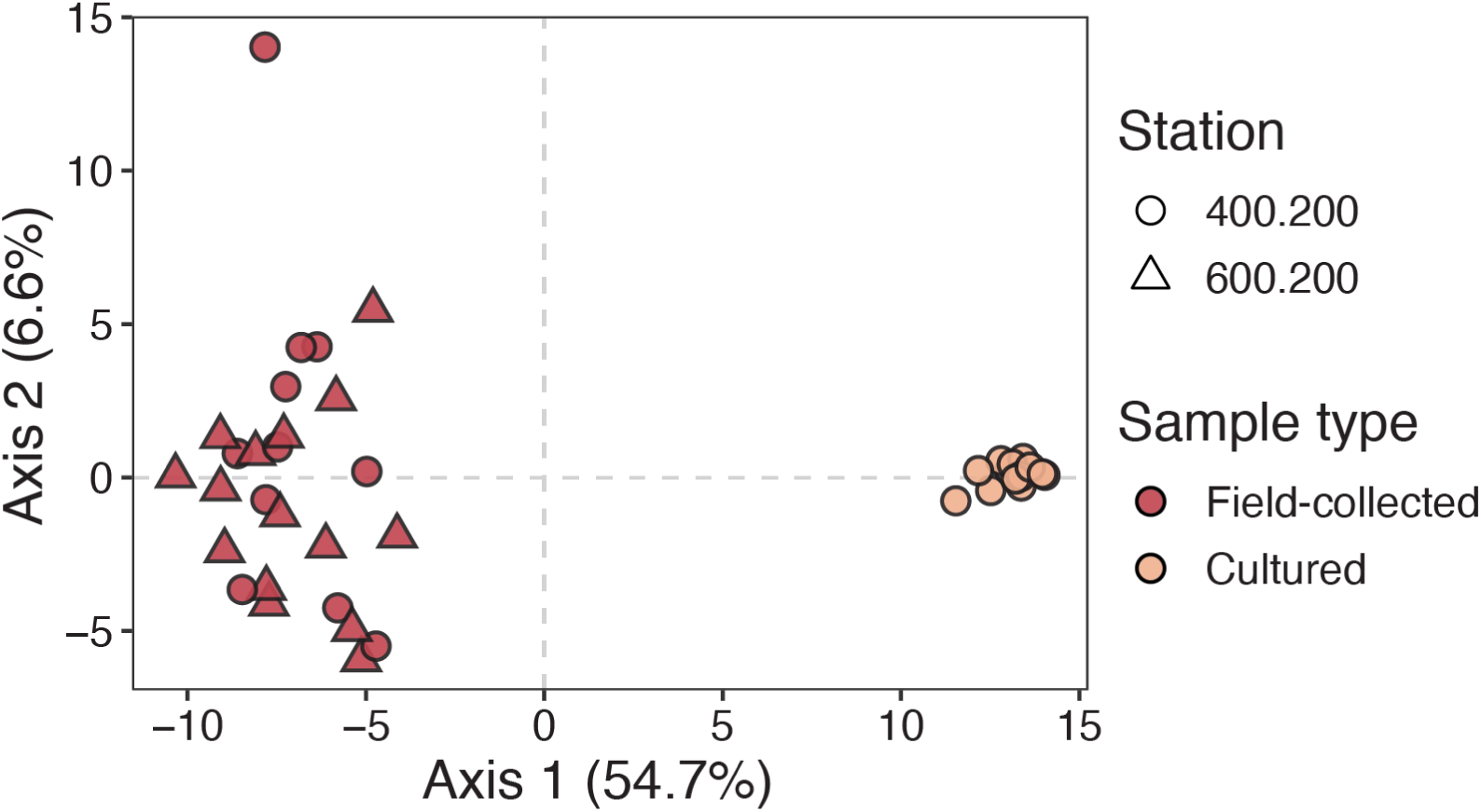
Principal Coordinates Analysis (PCoA) of Aitchison distances between bacterial communities associated with individual field-collected and cultured *Phaeocystis antarctica* colonies. Point shape represents PAL LTER grid station of origin (either directly collected from or used to start a culture) and point color represents sample type (field-collected or cultured). Two main clusters are visible: communities associated with field-collected colonies and communities associated with cultured colonies. These clusters separate along the primary axis (x-axis), which explains 54.7% of the variability. The secondary axis explains only 6.6% of the variability and while there is some spread among field-collected colony communities on this axis, there is no separation by station.

**Supplemental Figure 5.**
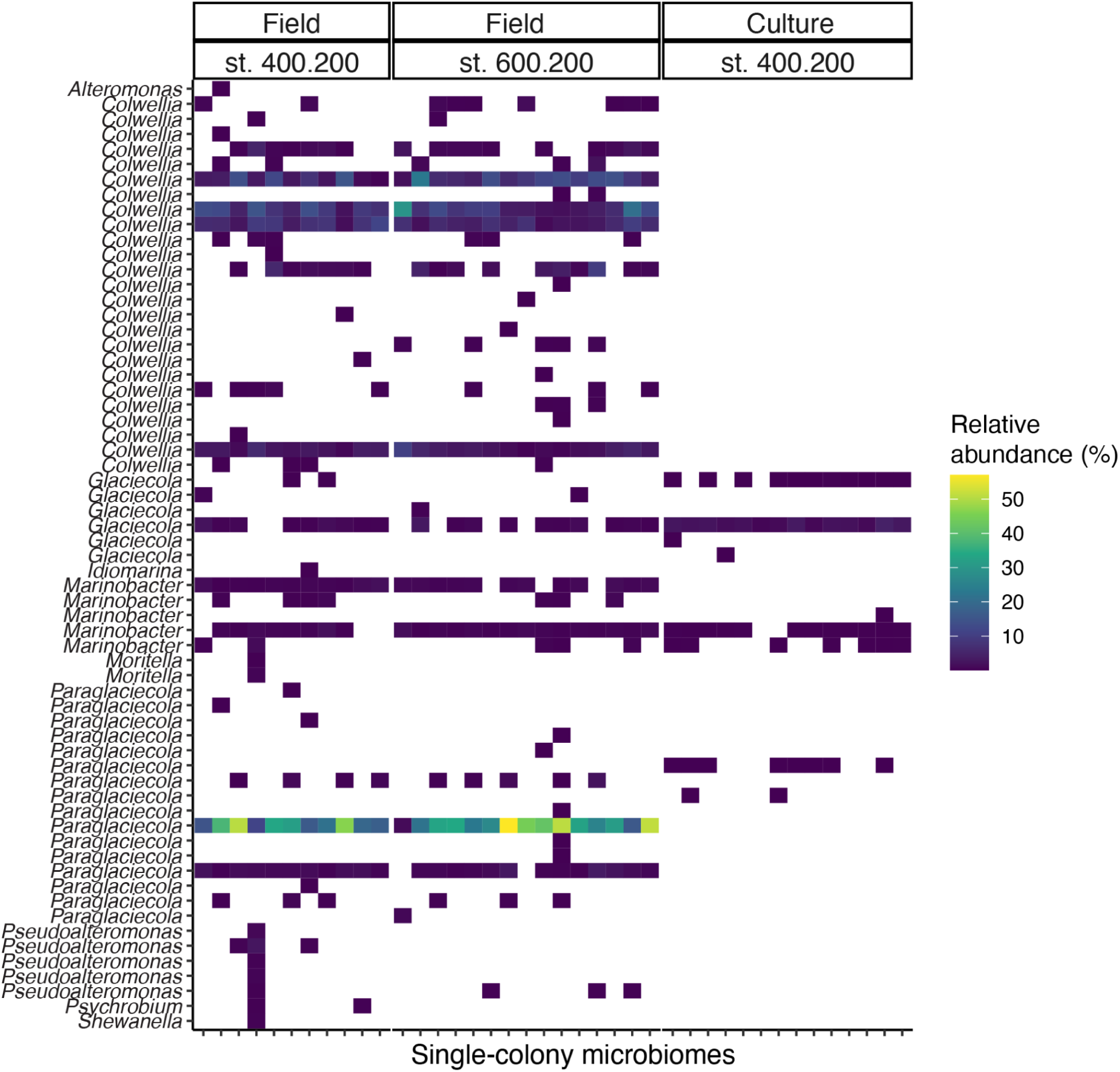
Relative abundance of Alteromonadales Amplicon Sequence Variants (ASVs) in field-collected and cultured colony microbiomes. Alteromonadales was the most dominant bacterial group in field-collected colonies but made up a small proportion of the cultured colony microbiomes. A single *Paraglaciecola sp.* ASV made up a large proportion of all field-collected colony microbiomes. Similarly, three *Colwellia sp*. ASVs were abundant in all field-collected colony microbiomes but were absent in cultured colony microbiomes. *Marinobacter* and *Glaciecola* ASVs were the only Alteromonadales ASVs detected in cultured colony microbiomes.

**Supplemental Figure 6.**
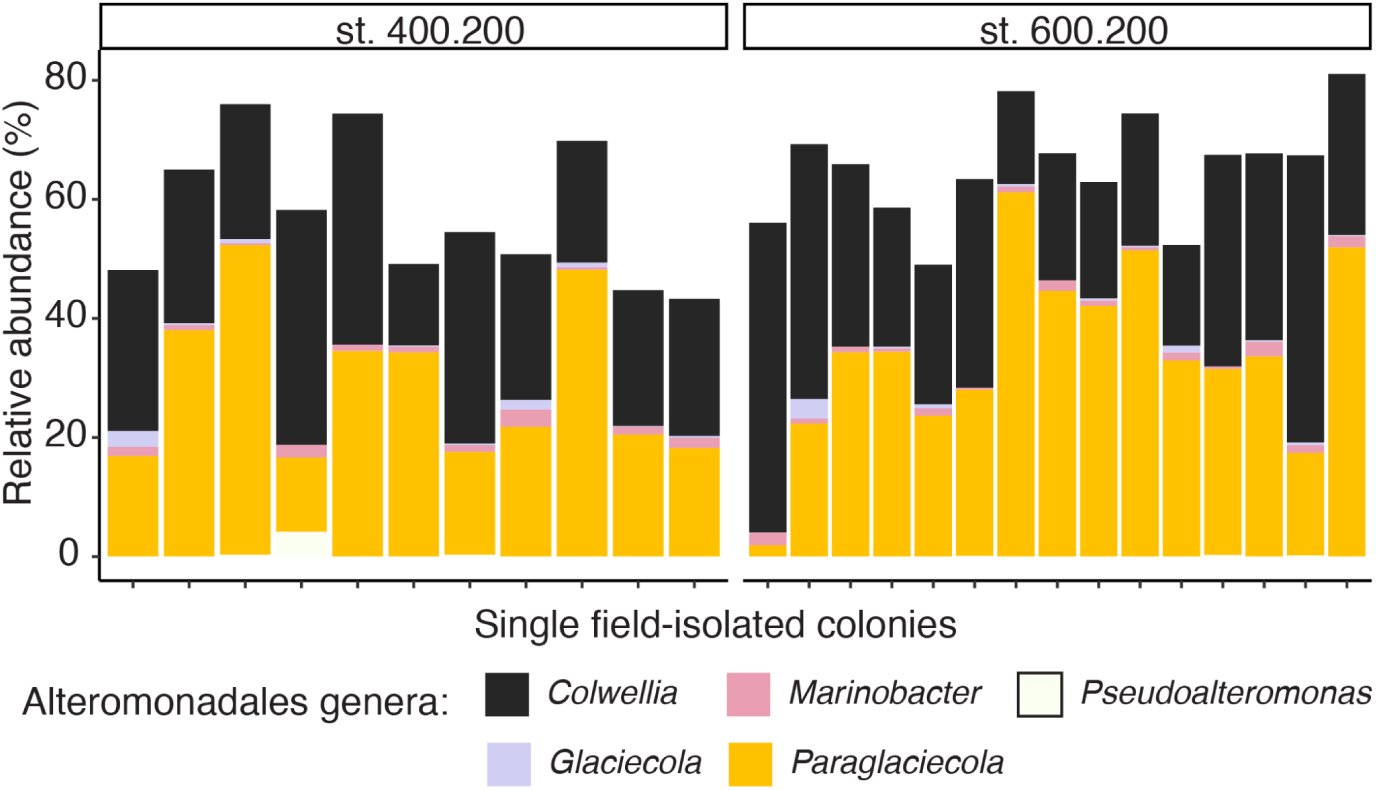
Relative abundance of Alteromonadales genera in field-collected colony microbiomes. The majority of Alteromonadales ASVs in field-collected colony microbiomes are *Colwellia sp*. or *Paraglaciecola sp.*, and these two genera contribute roughly similar proportions of sequences across the colony replicates.

**Supplemental Figure 7.**
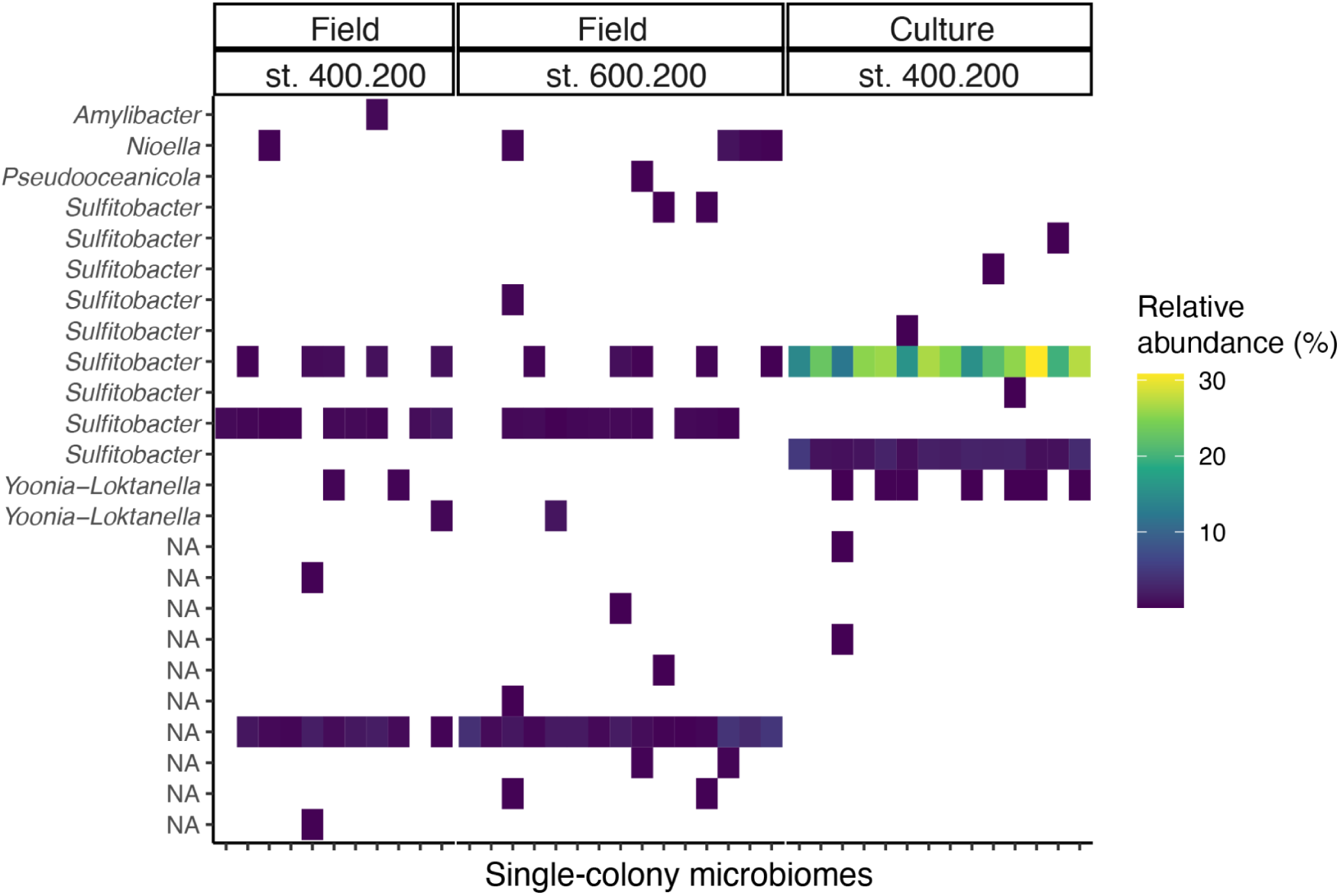
Relative abundance of Rhodobacterales Amplicon Sequence Variants (ASVs) in field-collected and cultured colony microbiomes. Rhodobacterales was the second most abundant group in cultured-colony microbiomes, but made up a small proportion of the field-collected colony microbiomes. A single *Sulfitobacter sp.* ASV made up a large proportion of all cultured colony microbiomes.

**Supplemental Figure 8.**
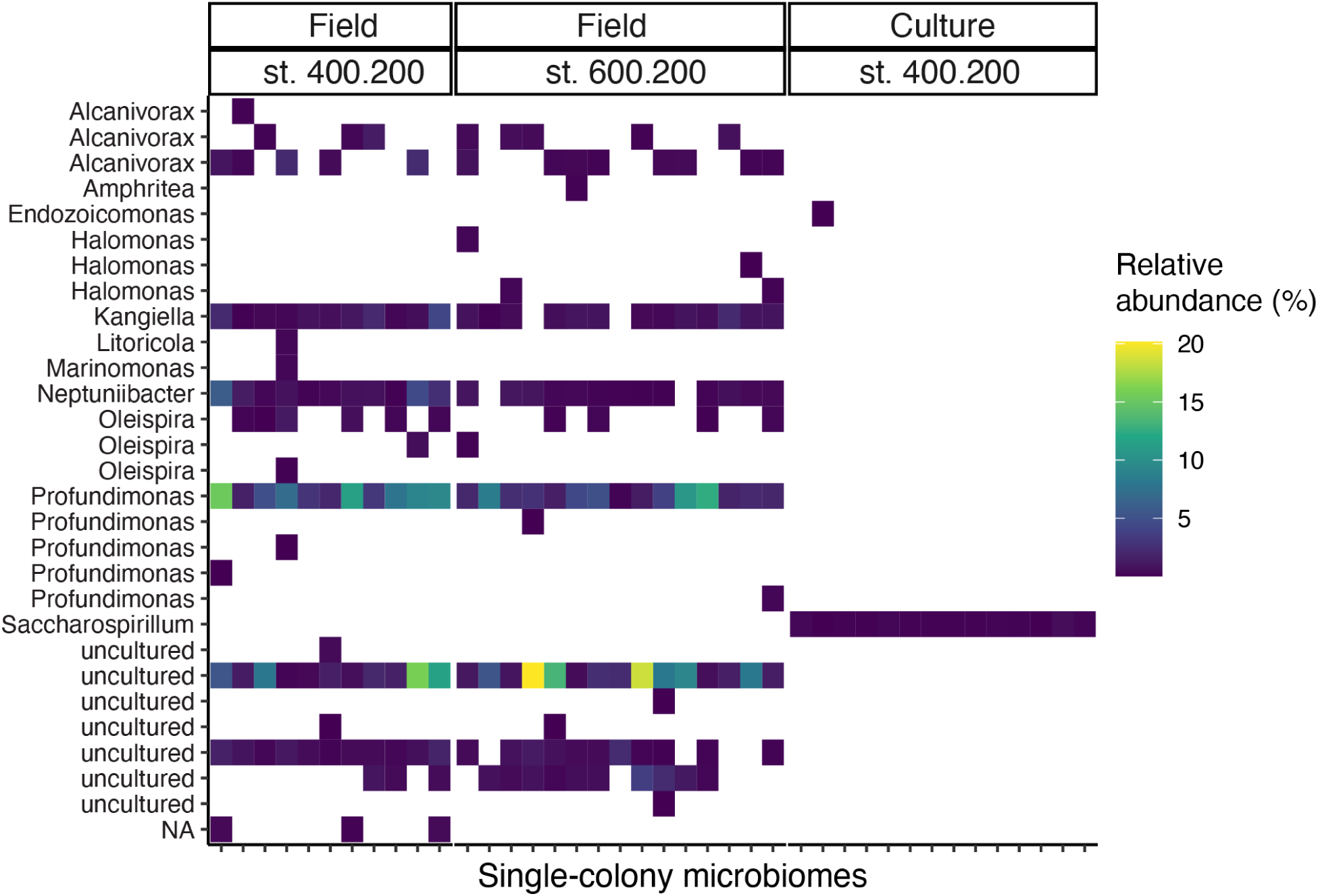
Relative abundance of Oceanospirillales Amplicon Sequence Variants (ASVs) in field-collected and cultured colony microbiomes. Alteromonadales was the second most dominant bacterial group in field-collected colonies but made up a small proportion of the cultured colony microbiomes. Two ASVs comprised a large proportion of all field-collected colony microbiomes, with one belonging to the genus Profundimonas (family: Nitrincolaceae) and the other being most similar to sequences from uncultured bacteria in the family Nitrincolaceae. Both of these ASVs were absent in cultured colony microbiomes. Instead, a *Saccharospirillum* ASV was the sole prevalent Oceanospirillales ASV in cultured colony microbiomes. *Saccharospirillum spp.* have been isolated from algal sediments and mangrove and seagrass rhizospheres, and while they can survive at low temperatures (4°C) their optimum growth temperatures are higher (25°C) (Yang et al. 2020).

**Supplemental Figure 9.**
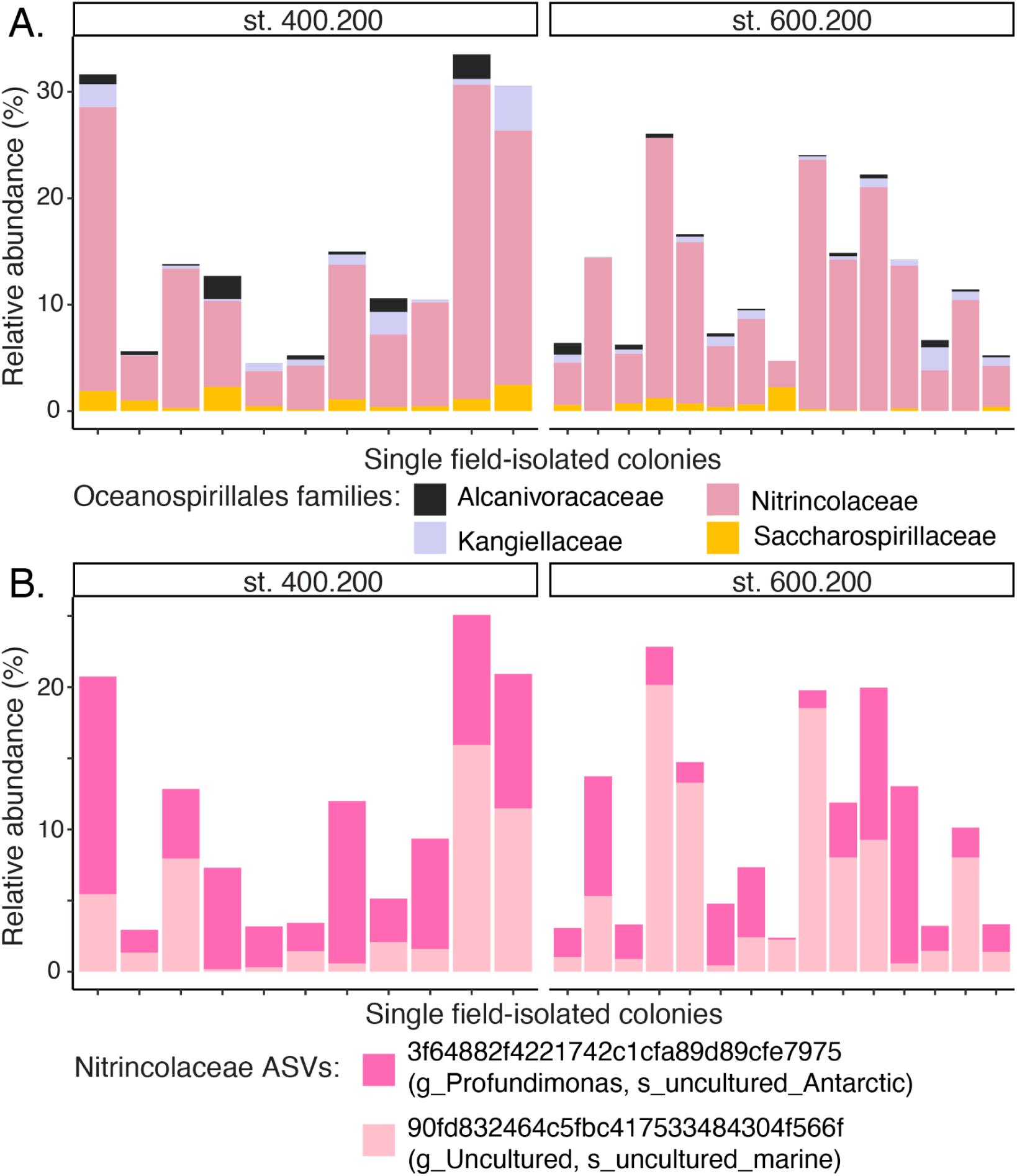
Relative abundance of Oceanospirillales families (A) and ASVs (B) in field-collected colony microbiomes. (A) Most Oceanospirillales ASVs in field-collected colony microbiomes belong to the family Nitrincolaceae. However, the family Saccharospirillaceae, which comprises all Oceanospirillales ASVs in cultured colony microbiomes, is detected in many field-collected colony microbiomes. (B) Just two ASVs comprise nearly the whole Nitrincolaceae contribution to field-collected colony microbiomes.

**Supplemental Figure 10.**
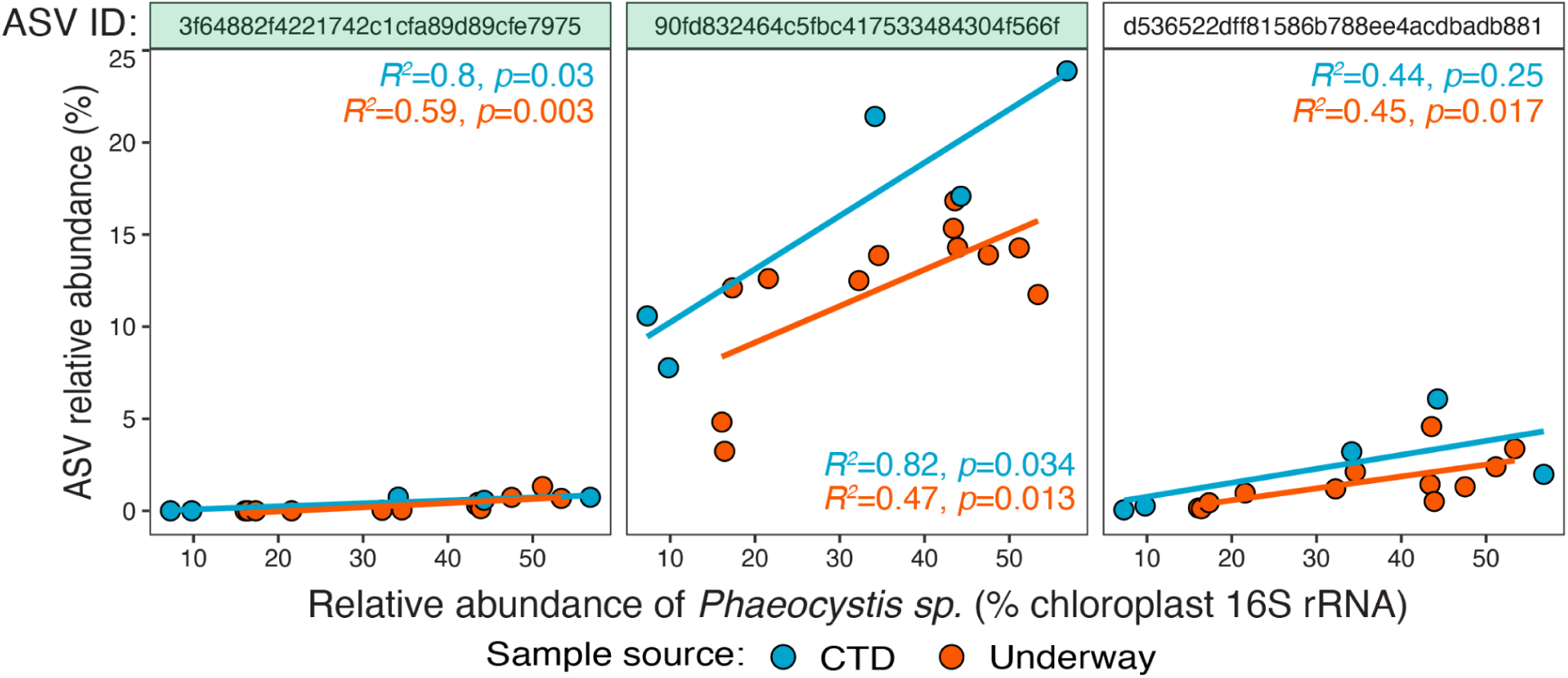
Relationship between the relative abundance of Oceanospirillales (family: Nitrincolaceae) and *Phaeocystis sp.* ASVs in surface seawater samples from the Palmer LTER grid. ASV IDs highlighted in green are the two most abundant Oceanospirillales ASVs in field-collected colony microbiomes. ASVs 90fd832464c5fbc417533484304f566f (center) and d536522dff81586b788ee4acdbadb881 (right) were the two most abundant Oceanospirillales ASVs in environmental samples (that were also found in field-collected colony microbiomes). All three ASVs were significantly correlated with *Phaeocystis* abundance in CTD (Niskin) and/or underway samples.

